# *Calotropis gigantea* extract induces apoptosis through extrinsic/intrinsic pathways and upregulation of reactive oxygen species in non-small-cell lung cancer cells

**DOI:** 10.1101/441162

**Authors:** Jiyon Lee, Huiju Jang, Yesol Bak, Jong-Woon Shin, Hang Jin, Yong-In Kim, Hyung Won Ryu, Sei Ryang Oh, Do-Young Yoon

**Author notes:** Corresponding author: (D.Y. Yoon).

## Abstract

**Background:** *Calotropis gigantea* (CG) plant grows in Asia and tropical Africa. However, the precise mechanisms of its anticancer effects have not yet been examined in human non-small cell lung cancer (NSCLC) cells, A549 and NCI-H1299 cells.

**Purpose:** This study was focused on the anti-cancer effects of CG extract on non-small cell lung cancer (NSCLC) cells.

**Methods:** The cytotoxic effects of CG extract on NSCLC, A549 and NCI-H1299 cells, were detected by MTS assay, microscope and DAPI staining. Apoptosis was determined by annexin V-FITC/PI staining, cell cycle analysis, western blotting, quantitative polymerase chain reaction, and JC-1 staining.

**Results:** First, CG showed significant dose-dependent cytotoxicity in NSCLC, A549, and NCI-H1299 cells. In addition to induction of caspase-8 processing, CG induced apoptosis by upregulating mRNA expression levels of extrinsic pathway molecules such as Fas, Fas ligand (FasL), Fas-associated protein with death domain (FADD) and death receptor 5 (DR5). Also, mitochondrial membrane potential (MMP) was collapsed, and intrinsic pathway molecules such as poly (ADP-ribose) polymerase (PARP), caspase-3, and caspase-9 were processed by CG. Moreover, reactive oxygen species (ROS) were generated in a CG dose-dependent manner, and inhibition of ROS by NAC, ROS scavenger, recovered A549 and NCI-H1299 cell viability.

**Conclusion:** These results indicate that CG causes apoptosis by activating the extrinsic and intrinsic pathways and generating ROS in NSCLC cells. These results suggest that CG can be used as a lung cancer therapeutic agent.

## Introduction

Lung cancer, also known as lung carcinoma, is one of the most common diseases in the world [1]. However, owing to very few therapies being available, diverse studies on lung cancer are needed. Lung cancer is classified into non-small cell lung cancers (NSCLCs) and small cell lung cancers (SCLCs) [2, 3]. SCLC is a neuroendocrine tumor type, and the size of cells in these cancers is smaller than those in NSCLC. In contrast, NSCLC includes squamous cell carcinomas, large cell carcinomas, and adenocarcinomas. Among these NSCLCs, A549 lung cancer cells (wild-type p53) [4] and p53 null NCI-H1299 lung cancer cells are human non-small cell lung carcinoma cell lines.

Lung cancer is caused by uncontrolled cell growth in lung tissues because of defects in cancer suppressor genes [5], which results in the failure of apoptotic signaling and thus cell proliferation. In other words, inducing programmed cell death significantly reduces cancer cell numbers [6]. Apoptosis is a process of programmed cell death that controls cell division. If cells are old, our body causes them to commit suicide by activating an intracellular death process. Cells die by two apoptotic pathways: the intrinsic pathway and extrinsic pathway. First, the intrinsic pathway of apoptosis starts when the mitochondria outer membrane becomes permeable in response to intracellular stressors such as DNA damage, growth factor impairment, or oncogene activation. Cytochrome c and pro-apoptosis proteins are released from permeable mitochondria in a Bax/Bak-dependent manner [7]. These releases are regulated by Bcl-2 family members such as Bcl-2 and Bcl-xL and BH-3 family members such as Bid. Bax and Bak are activated by members of the BH-3 family [8] and inhibited by members of the Bcl-2 family. Through the association of these proteins, cytochrome c forms a complex, the apoptosome [9], which is composed of cytochrome c and caspase-9. Caspase-9 is activated by the apoptosome, and cleaved caspase-9 promotes activation of caspase-3 and inactivation of poly (ADP-ribose) polymerase (PARP), whose function is DNA damage repair [10]. Through this mechanism, this pathway induces apoptosis of cells. Second, two direct mechanisms of extrinsic pathway initiation have been described. One is the tumor necrosis factor (TNF)-induced model, and the other is the Fas-FasL-mediated model. Ligand binding with the receptor recruits adaptor proteins such as FADD and TRADD [11] and initiators such as pro-caspase-8 and pro-caspase-10. These complexes activate caspase-8 and caspase-10. Cleaved caspase-8 induces apoptosis by cleaving caspase-3. In addition, the extrinsic and intrinsic pathways are associated with caspase 8, which cleaves Bid into truncated Bid (tBid), the activated form of the protein. tBid induces the Bax/Bak-dependent release of cytochrome c. Finally, cleaved caspase-3 and inactivated PARP are triggered through both apoptotic pathways to dismantle the cells [12].

Programmed cell death can be initiated by various types of stress-induced damage. Reaction oxygen species (ROS) production [13] is a critical stressor that causes cell death, especially through the induction of apoptosis. Several reports have shown that ROS plays in important role in the apoptosis of cancer cells. The production of ROS such as ^•^OH, O_2_−, and H_2_O_2_, caused by various external stimuli, is related to the inhibition of cell proliferation [14]. Low doses of H_2_O_2_ produce ^•^OH radicals, and an alternative oxidant/antioxidant pathway induces apoptosis. Furthermore, Bcl-2 controls not only apoptotic death but also the antioxidant pathway [15]. Cytochrome c release is suppressed by Bcl-2, and superoxide production is prevented when Bcl-2 is overexpressed [16]. In the cascade, a higher oxidant concentration decreases cell proliferation and finally induces apoptosis.

*Calotropis gigantea* (CG) is a tall and waxy flower that is mainly distributed in Cambodia, Indonesia, Malaysia, the Philippines, Thailand, Sri Lanka, India, China, Pakistan, Nepal, and tropical Africa. A hypoglycemic effect of the extract of flowers and leaves of CG [17] has been shown, with serum glucose levels that decreased with CG extract treatment. However, the presence of an anti-cancer effect of CG extract on lung cancer cells has not been assessed. Here, we report the presence of apoptosis signaling when lung cancer cells were treated with CG extract. Our results indicate a potential anti-cancer effect of CG extract on lung cancer cells.

## Materials and Methods

### Reagents and antibodies

A stock solution of CG (30 mg/mL) in dimethyl sulfoxide (DMSO) was stored in the dark at −20°C and diluted in Rowell Park Memorial Institute (RPMI)-1640 medium immediately before use. CellTiter 96^®^ AQueous One Solution Cell Proliferation Assay Reagent [MTS, 3-(4,5-dimethylthiazol-2-yl)-5-(3-carboxymethoxyphenyl)-2-(4-sulfophenyl)-2H-tetrazolium] was purchased from Promega (Madison, WI, USA), and propidium iodide (PI) was from Sigma-Aldrich (St. Louis, MO, USA). Antibodies specific to PARP, caspase-3, caspase-8, caspase-9, Bcl-2, Bcl-xL, Bax, Bid, and cytochrome c were from Cell Signaling Technology (Beverly, MA, USA). The antirabbit IgG horseradish peroxidase (HRP)-conjugated secondary antibody and anti-mouse IgG HRP-conjugated secondary antibody were from Millipore (Billerica, MA, USA). Antibodies specific to p21, p27, cyclin D1, cyclin E, cyclin A, and glyceraldehyde 3-phosphate dehydrogenase (GAPDH) were purchased from Santa Cruz Biotechnology (Santa Cruz, CA, USA). JC-1 (5,50,6,60-tetrachloro-1,10,3,30-tetraethyl benzimidazoly carbocyanine chloride) was obtained from Enzo (New York, USA), and FITC-annexin V apoptosis detection kit I was purchased from BD Biosciences (San Diego, CA, USA). DCF-DA was procured from Abcam (Cambridge, UK).

### Plant material and preparation

Ethanol extract of whole plant of *C. gigantea* (L.) W.T. Aiton (Asclepiadaceae) was supplied by Foreign Plant Extract Bank (no. FBM085-042; Daejeon, Korea). The plant was collected in Yunnan Province of China in 2008 and authenticated by Jin Hang, The Chief of Medicinal Plants Research Institute, Yunnan Academy of Agricultural Sciences (YAAS) (Yunnan, China). A voucher specimen (YASS3533-2) is kept at the herbarium of YAAS. In preparation steps, the air-dried whole plant of *C. gigantea* sample (100.0 g) was mixed in 95% ethanol (800 mL × 2), and the mixture was shaken at room temperature for 2 h. The extracts were combined and concentrated in vacuo at 40 °C to produce a dried extract, which was used for phytochemical analysis and biological activities.

### UPLC-QTof-MS analysis

Tentative identification of compounds from *C. gigantea* extracts were carried out using ACQUITY UPLC (Waters Corporation, Milford, MA) system connected to Micromass QTof Premier™ mass spectrometry (Waters Corporation, Milford, MA) with an electrospray ionization device, operating in the negative ion mode, using the following operation parameters: The operation parameters in the negative ion mode were capillary voltage, 2,300 V; cone voltage, 50 V; ion source temperature, 110°C; desolvation temperature, 350 °C; flow rate of desolvation gas (N_2_), 500 L/h; mass scan range, 100–1500 Da; scan time, 0.25 s. Leucine enkephalin was used as the reference compound (*m/z* 554.2615 in negative ion mode). The gradient elution program was as follows: 0 min, 10% B; 0–1.0 min, 10% B; 1.0–12.0 min, 10–100% B; wash for 13.4 min with 100% B; and a 1.6 min recycle time. The injection volumes were 2.0 μL, and the flow rate was 0.4 mL/min.

### Cell culture

A549 and NCI-H1299 cells were purchased from the American Type Culture Collection (ATCC: Manassas, VA, USA). Cells were cultured in RPMI 1640 medium (Welgene, Gyeongsan-si, South Korea) containing 10% (v/v) heat-inactivated fetal bovine serum (FBS: Hyclone Laboratories, Logan, UT, USA). Cells were incubated at 37°C in an atmosphere of 5% CO_2_/95% air with saturated humidity.

### Cell viability assays

Cell viability was examined with 3-(4,5-dimethylthiazol-2-yl)-5-(3-carboxymethoxyphenyl)-2-(4-sulfophenyl)-2H-tetrazolium (MTS) assay. A549 cells were seeded at 0.7 × 10^4^ cells and NCI-H1299 cells at 0.9 × 10^4^cells in 100 μL medium/well in 96-well plates for overnight growth. After 24 h, various concentrations of CG extract were added, and the cells were incubated for an additional 24 and 48 h. The medium (100 μL) was removed and incubated with 100 μL of MTS with PMS mix solution for 40 min to 1 h at 37°C. The optical densities in each well were measured at 492 nm with an ELISA reader (Apollo LB 9110, Berthold Technologies GmbH, Zug, Switzerland).

### DAPI staining

A549 cells (0.5 × 10^4^ cells) and NCI-H1299 cells (0.7 × 10^4^ cells) were seeded in each well of slide plate and incubated overnight. After that, cells were treated with various concentrations of CG. After 48 h, plates were washed with PBS and cells were fixed with 4 *%* paraformaldehyde and stained with DAPI staining solution for 10 seconds. The plates were washed with PBS and completely dried. Plates were mounted with cover slip. The stained cells were observed using Fluorescence microscope (Olymous, Tokyo, Japan).

### Annexin V/PI staining

A549 cells (1.5 × 10^5^ cells) and NCI-H1299 cells (2.0 × 10^5^ cells) were seeded in 1.5 mL medium/well in 6-well plates for overnight growth. Cells were treated with various concentrations of CG extract for 48 h, harvested using trypsin, and washed with PBS. Annexin V and PI staining were performed using a FITC-Annexin V Apoptosis Detection Kit I (BD Biosciences, San Jose, CA, USA) according to the manufacturer’s instructions. Data were analyzed using flow cytometry, employing a FACSCalibur instrument and CellQuest software (BD Biosciences, San Jose, CA, USA).

### Cell cycle analysis

The cell cycle distribution was analyzed by PI (propidium iodide) staining and flow cytometry. A549 (1.5 × 10^5^ cells) and NCI-H1299 cells (2.0 × 10^5^ cells) were seeded in 1.5 mL medium/well in 6-well plates for overnight growth and treated with various concentrations of CG extract. After 48 h, cells were harvested with trypsin and fixed with 80% ethanol for > 1 h. Next, the cells were washed twice with cold phosphate-buffered solution (PBS) and centrifuged, and the supernatants were removed. The pellets were re-suspended and stained in PBS containing 50 μg/mL PI and 100 μg/mL RNase A for 20 min in the dark. DNA content was analyzed by flow cytometry using a FACSCalibur instrument and CellQuest software (BD Biosciences, San Jose, CA, USA).

### Real-time quantitative polymerase chain reaction (qPCR)

A549 cells treated with CG for 48 h were harvested. The harvested cells were lysed in 1 mL easy-BLUE™ (iNtRon Biotechnology, SungNam, Korea). The RNA was isolated according to the manufacturer’s instructions. cDNA products were obtained using M-MuL V reverse transcriptase (New England Biolabs, Beverly, MA, USA). Real-time qPCR was performed using a relative quantification protocol using Roter-Gene 6000 series software 1.7 (Qiagen, Venlo, Netherlands) and a SensiFAST™ SYBR NO-ROX Kit (BIOLINE, London, UK). Each sample contained one of the following primer sets: *FAS* F: 5′-CGGACCCAGAATACCAAGTG-3′ and R: 5′-GCCACC CCAAGTTAGATCTG-3′; *FASL F:* 5′-GGGGATGTT TCAGCTCTTCC-3′ and R: 5′-GTGGCCTAT TTG CTT CTCCA-3′; *DR5* F: 5′-CACCTTGTACACGATGC TGA-3′ and R: 5′-GCTCAACAA GTGGTCCTCAA-3′; *FADD* F: 5′-GGGGAAAGATTGGAGAAGGC-3′ and R: 5′-CAGTTCTCAGTGA CTCCCG-3′; *SOD2* F: 5′-TATAGAAAGCCGAGTGTTTCCC-3′ and R: 5′-GGGATGCCTTTC TAGTCCTATTC-3′: *catalase* F: 5′-GGG ATCTTTTAACGCCATT-3′ and R: 5′-CCAGTTTACCAACTGGATG-3′; *thioredoxin* F: 5′-GAAGCTC TGT TTGGTGCTTTG-3′ and R: 5′-CTCGATCTGCTTCACCATCTT-3′ *GAPDH* F: 5′-GGCTGCTTTTAAC TCT GGTA-3′ and R: 5′-TGGAAGATGGTGATGGGATT-3′.

### Western blot analysis

A549 cells and NCI-H1299 cells were treated with CG at various concentrations for 48 h, harvested, and then washed with PBS. After centrifugation (13,000 rpm, 1 min, 4°C), the cell pellets were re-suspended in a lysis buffer containing 50 mM Tris (pH 7.4), 1.5 M sodium chloride, 1 mM EDTA, 1% NP-40, 0.25% sodium deoxycholate, 0.1% sodium dodecyl sulfate (SDS), and a protease inhibitor cocktail. The cell lysates were vortexed by a rotator at 4°C for 1 h and clarified by centrifugation at 13,000 rpm for 30 min at 4°C. The protein content was estimated using a Bradford assay (Bio-Rad Laboratories, Hercules, CA, USA) and UV spectrophotometer. The cell lysates were loaded onto a 10–12 *%* gel and separated by SDS-polyacrylamide gel electrophoresis (PAGE). The protein bands were transferred to polyvinylidene difluoride (PVDF) membranes (Millipore, Billerica, MA, USA). Next, the membranes were blocked with Tris-buffered saline containing Tween-20 (TBST) (2.7 M NaCl, 1M Tris-HCL, 53.65 mM KCL, and 0.1% Tween-20, pH 7.4) and 5% skim milk for 30 min at room temperature. The membranes were incubated overnight at 4°C with primary antibodies targeting specific proteins. After three washes with TBST for 10 min each, the membranes were incubated with a secondary antibody (HRP-conjugated anti-rabbit or anti-mouse IgG) for 2 h at room temperature. After three washes with TBST, the blots were analyzed using West-Zol and a western blot detection system (iNtRON Biotechnology, South Korea).

### Nuclear and cytoplasmic fractionation

A549 cells and NCI-H1299 cells treated with CG were collected and fractionated using NE-PER nuclear and cytoplasmic extraction reagents (Thermo Fisher Scientific Inc., USA) according to the manufacturer’s instructions. The treated cells were harvested with trypsin-EDTA and centrifuged at 500 × *g* for 3 min. The cell pellets were suspended in 200μL cytoplasmic extraction reagent I by vortexing. The suspensions were incubated on ice for 10 min, followed by the addition of 11 μL second cytoplasmic extraction reagent II. These reaction mixtures were vortexed for 5 s, incubated in iced for 1 min, and centrifuged at 16,000 × *g* for 5 min. Then, the supernatant fractions were transferred to pre-chilled tubes. The insoluble fractions were resuspended in 100 μL nuclear extraction reagent by vortexing for 15 s. The reaction tubes were incubated on ice for 10 min and centrifuged for 10 min at 16,000 × *g*. The resulting supernatant constituting the nuclear extract was used for subsequent experiments.

### Analysis of mitochondrial membrane potential (MMP)

We evaluated MMP (ΔΨm) by JC-1 staining and flow cytometry. A549 (3.8 × 10^5^ cells) and NCI-H1299 (4.3 × 10^5^ cells) cells were seeded into 3 mL medium/60 mm culture dishes. The cells were treated with various concentrations of CG. Cells were harvested with trypsin-EDTA and transferred into 1.5 mL tubes. JC-1 (5 μg/mL) was added to the cells and mixed until it was completely dissolved, followed by incubation of cells in the dark for 10 min at 37°C in an incubator. The cells were centrifuged (300 × *g*, 5 min, 4°C), washed twice with PBS, and resuspended in 200 μL PBS. The solutions were analyzed using a FACSCalibur instrument and Cell Quest software, in minimal light.

### Detection of intracellular levels of ROS

We used a DCF-DA cellular ROS detection assay kit (Abcam, UK) to detect the accumulation of intracellular ROS in A549 and NCI-H1299 cells. A549 (0.7 × 10^4^cells) and NCI-H1299 (0.9 × 10^4^cells) cells were seeded into dark 96-well plates and incubated 24 h. The cells were then stained with 25 μM 2′,7′-dichlorofluorescin diacetate (DCF-DA) for 45 min and treated with various concentrations of CG (0, 3.75, 7.5, and 15 μg/mL) for 48 h. The fluorescence intensity of each well was quantified with a fluorescence microplate reader (Gemini EM, Molecular Devices, USA) at excitation/emission = 485/538 nm. The experiments were performed at least three times.

## Results

### Tentative identification of phytochemicals in CG

UPLC-PDA-QTof-MS analyses were carried out using a C18 column with a linear gradient of acetonitrile/water (Fig. 1). All peaks were characterized using Mass. Table 1 shows the retention times, UV-Vis absorption maxima, mass spectral data of molecular ions of these compounds in the CG extract, namely quercetin 3-rutinoside, kaempferol-4′-O-rutinoside, kaempferol-3-O-rutinoside, isorhamnetin-3-O-rutinoside, deglucoerycordin, 15ß-hydroxycalotropin, frugoside, and trihydroxy octadecenoic acid. Various rutinosides and a high level of isorhamnetin-3-O-rutinoside, particularly, were found in the CG extract.

**Figure 1.**
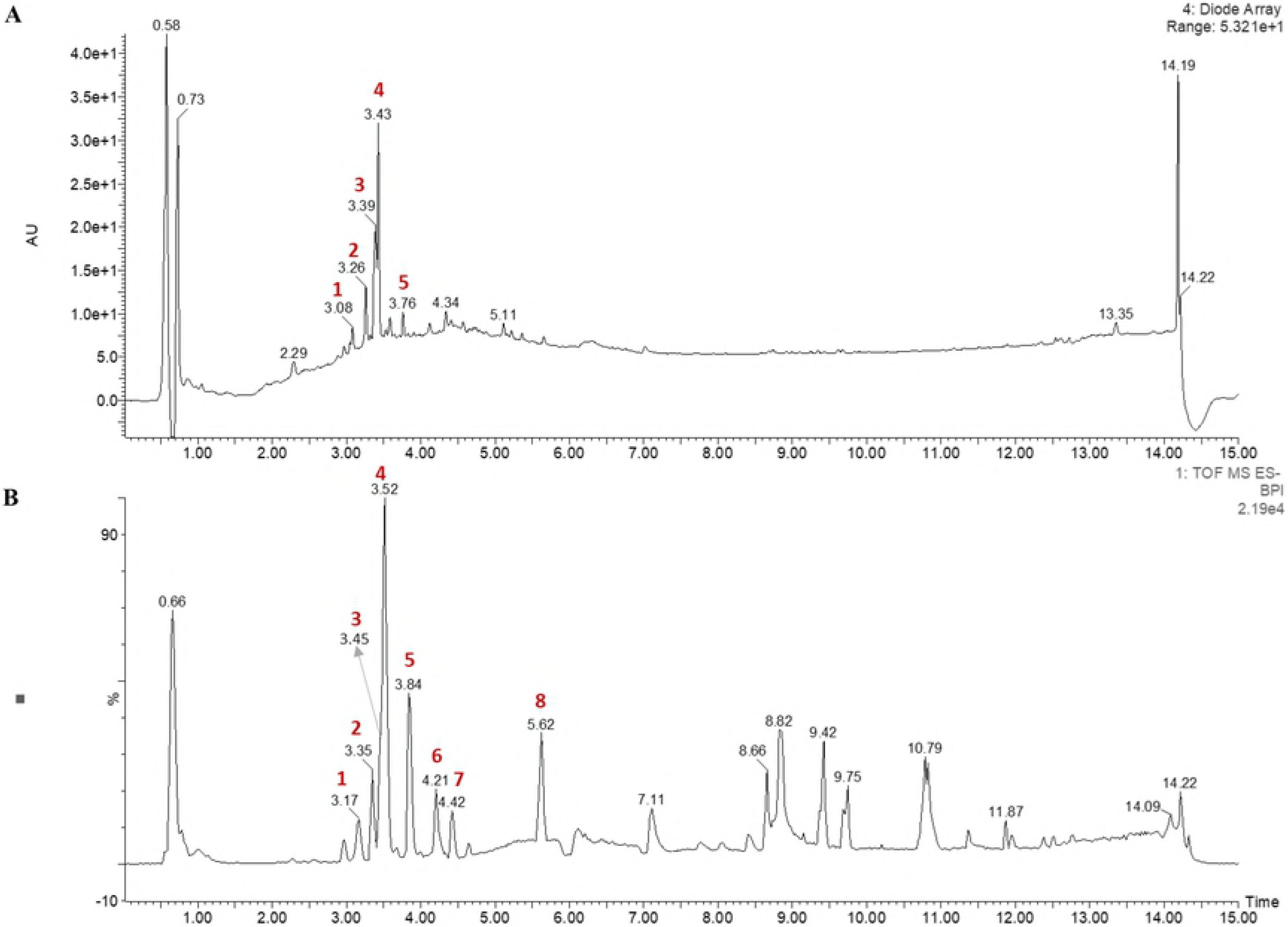
Representative UPLC-QTof-MS chromatograms of the methanol extracts from CG. Peak numbers refer to Table 1. The composition of CG was analyzed by UPLC-QTof-MS. Among the compounds in CG, isorhamnetin-3-O-rutinoside was a major component.

**Table 1.**
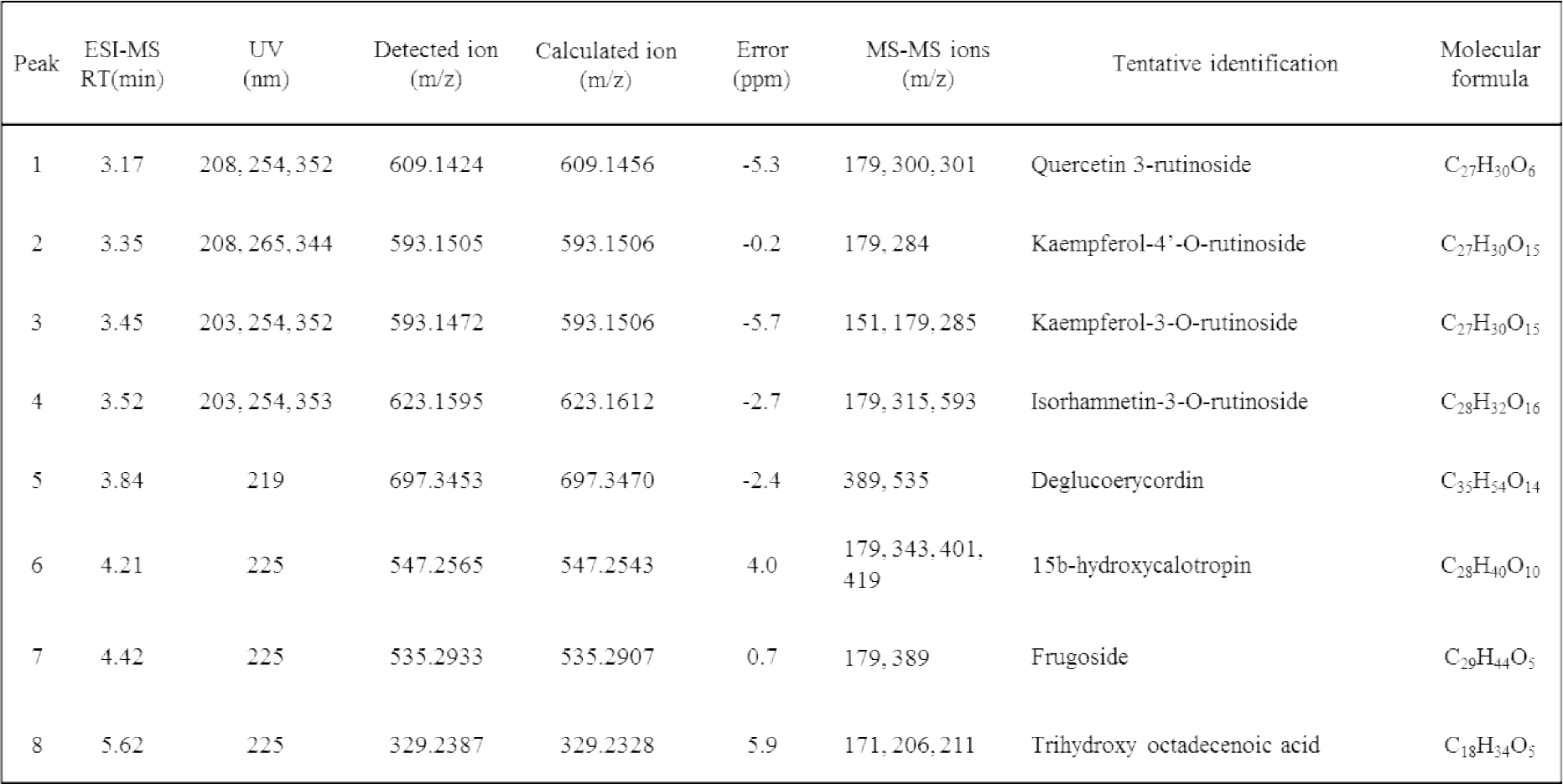
Quantitative HPLC analyses of composition in CG extraction.

### CG induces cytotoxic effects in human NSCLC cells

Several studies have been reported on the effects of plant extracts inhibiting tumor growth, metastasis, and angiogenesis [18]. This study was aimed to determine whether CG could induce cytotoxicity, examined in A549 and NCI-H1299 cells following treatment with various concentrations (up to 15 μg/ml for 24 and 48 h) of CG. The MTS assay showed that the cell viability rate declined in a dose-dependent manner by CG (Fig. 2A and B). The viabilities of A549 and NCI-H1299 cells treated with the highest (up to 15 μg/ml) CG concentrations at 48 h were both less than 50% that in DMSO (0.01%) treated control cells; the viabilities of A549 and NCI-H1299 cells treated with CG decreased significantly after 48 h. Therefore, CG extract was found to exert cytotoxic effects on lung cancer cells. Thus, our next study was focused on verifying the mechanism underlying NSCLC apoptosis following up to 15 μg/ml of CG for 48 h on NSCLC cells.

**Figure 2.**
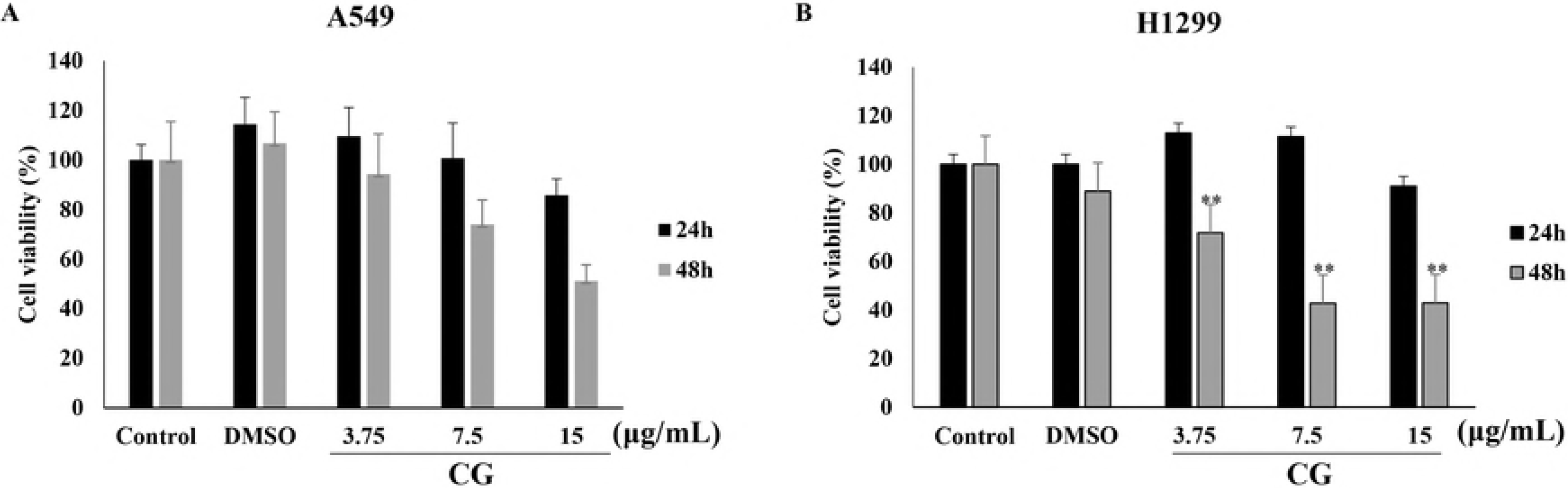
Cytotoxic effects of CG extract on human NSCLC cell lines. The viabilities of A549 (A) and NCI-H1299 cells (B). A549 and NCI-H1299 cells were treated for 24 and 48 h with CG extract. The untreated cells were compared with CG-treated cells. Results with \**p* < 0.05 and \*\**p* < 0.005 were considered statistically significant. Viability was analyzed by MTS assay.

### CG induces apoptosis in A549 and NCI-H1299 lung cancer cells

As the viabilities of A549 and NCI-H1299 cells treated with CG decreased dose dependently, cell morphological changes were observed by phase-contrast microscopy. As expected, these morphological changes were dependent on the dose of CG extract used to treat cells for 48 h. A549 (Fig. 3A) and NCI-H1299 (Fig. 3B) cells morphologies became more rounded and less interactions with surrounding cells were observed at high concentrations of CG than in DMSO (0.01%) treated control cells. This indicated that CG can change cell morphology, subsequently inducing cell death [19]. DAPI staining and fluorescence microscopy showed that CG-treated A549 cells (Fig. 3C) and NCI-H1299 cells (Fig. 3D) had condensed and their nuclei had fragmented—a marker of apoptosis—relative to the control cells. For further evidence of the effects of CG, CG treated A549 and NCI-H1299 cells were examined by Annexin V and PI staining [20]. When apoptosis occurs in the cells, lipid phosphatidylserine (PS) in the membranes of cells is translocated from the inner to the outer membrane, a so-called flip-flop movement, and Annexin V staining targets the PS [20]. Furthermore, the cell membranes had pores that facilitated apoptosis mediated via PI binding to DNA. The Annexin V-FITC/ PI staining indicated apoptosis in A549 (Fig. 3E) and NCI-H1299 cells (Fig. 3F) following treatment with CG extract. Cells of both types treated with CG for 48 h revealed increasing numbers of early and late apoptotic cells, whereas the number of live cells decreased. In A549 cells, the number of late apoptotic cells increased up to 35.30% with 15 μg/mL of CG treatment compared to that in control cells, 6.75 %. The rate of increase (3.42 % to 13.07 %) of late apoptotic cells among all NCI-H1299 cells was lesser than that of A549 cells. These results indicated that the death of A549 and NCI-H1299 cells induced by CG extract was thus mediated by apoptosis.

**Figure 3.**
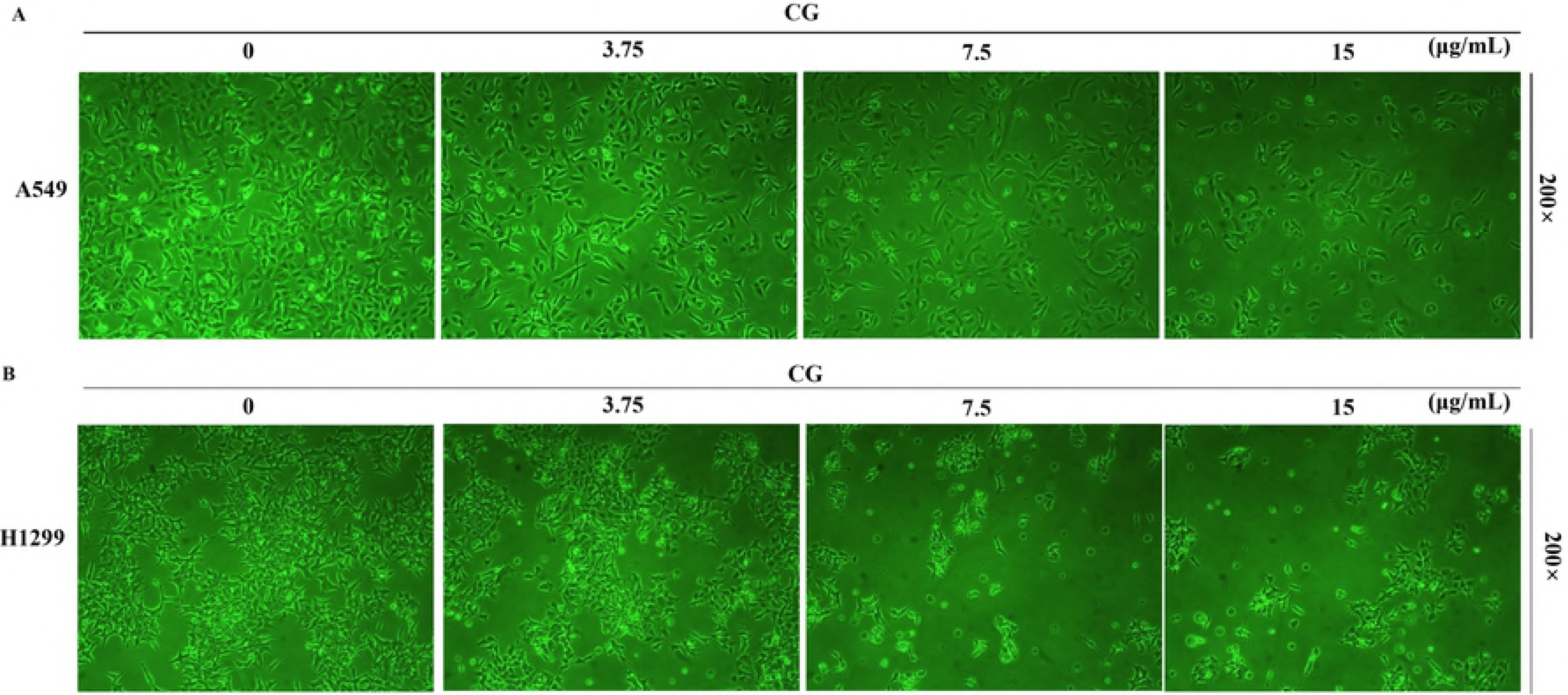
Effects of CG on viability and apoptosis in A549 and NCI-H1299 lung cancer cells. Microscopic images of A549 (A) and NCI-H1299 cells (B) treated with CG for 48 h. Fluorescence microscopic images showed that nuclear condensation and chromatin shrinkage were apparent in A549 (C) and NCI-H1299 cells (D) treated with CG for 48 h. After treatment with the indicated concentrations of CG for 48 h, A549 (E) and NCI-H1299 cells (F) were stained with Annexin V-FITC/PI. Untreated cells were compared with CG-treated cells.

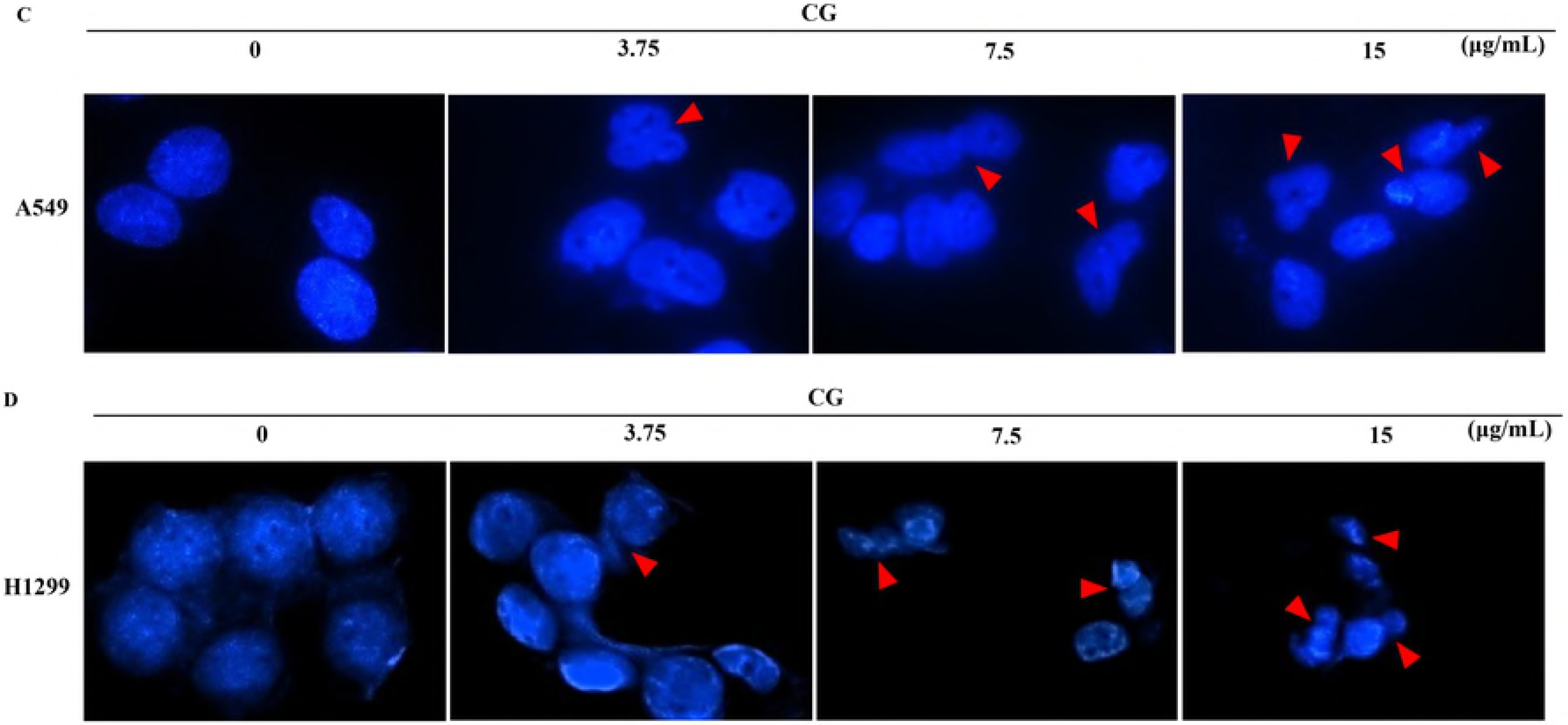

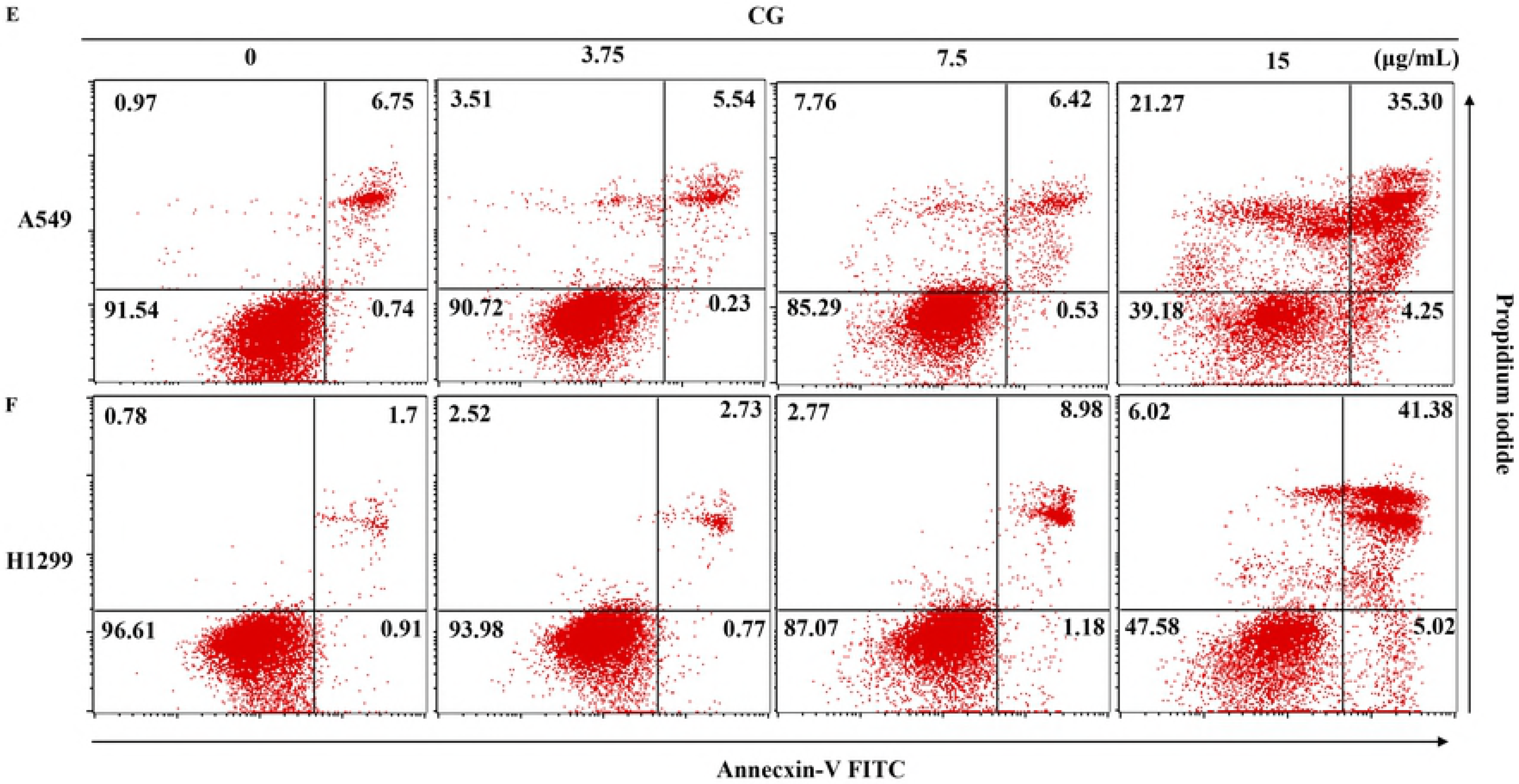

### CG modulates cell cycle-related factors in A549 cells and controls cell cycle progression in A549 and NCI-H1299 cells

Cell cycle disruption is a major cause of apoptosis in cancer calls [21], and many factors such as p53, p27, p21, and cyclins control cell cycle phases. p53 is known as a tumor suppressor gene [22]. When DNA has been damaged, it is activated to induce apoptosis. In addition, phosphorylated p53, p27 and p21, downstream factors of p53, regulate cyclin dependent kinase. The levels of p53 in A549 cells following treatment with 7.5 μg/mL and 15 μg/mL of CG extract were increased compared to that of the control, and 3.75 (μg/mL) concentrations and p27 was upregulated by CG as same as p53, but p21 was not (Fig. 4A). This suggests that p53 and p27 were stimulated by CG and induced the death of A549 cells by terminating the cell cycle. However, in NCI-H1299 p53-knockout cells, the cell cycle regulatory factors p27 and p21 were not affected by CG treatment (Fig. 4B). The cell cycle analyses of CG treated A549 (Fig. 4C) and NCI-H1299 cells (Fig. 4D) were performed by flow cytometry. Cells in the sub-G1 phase showed fragmented DNA, which is a marker of apoptosis [23, 24]. In our study, the cell cycle analysis showed that cells in the sub-G1 phase and S phase increased in a dose-dependent manner with CG treatment for A549 cells (Fig. 4E) and NCI-H1299 cells (Fig. 4F), whereas cells in the G0/G1 and G2/M phases decreased for both cell line. Furthermore, the cell cycle related factors cyclin D1 related to the sub-G1 phase and cyclin A related to the S phase, were downregulated as expected, but cyclin E in A549 cells was not altered by CG treatment (Fig. 4G). However, the levels of cyclin D1, E, and A decreased with treatment of NCI-H1299 cells (Fig. 4H). These results indicated that CG extract had negative effects on the A549 and NCI-H1299 cell cycles, reducing the restrictions against unlimited cell growth.

**Figure 4.**
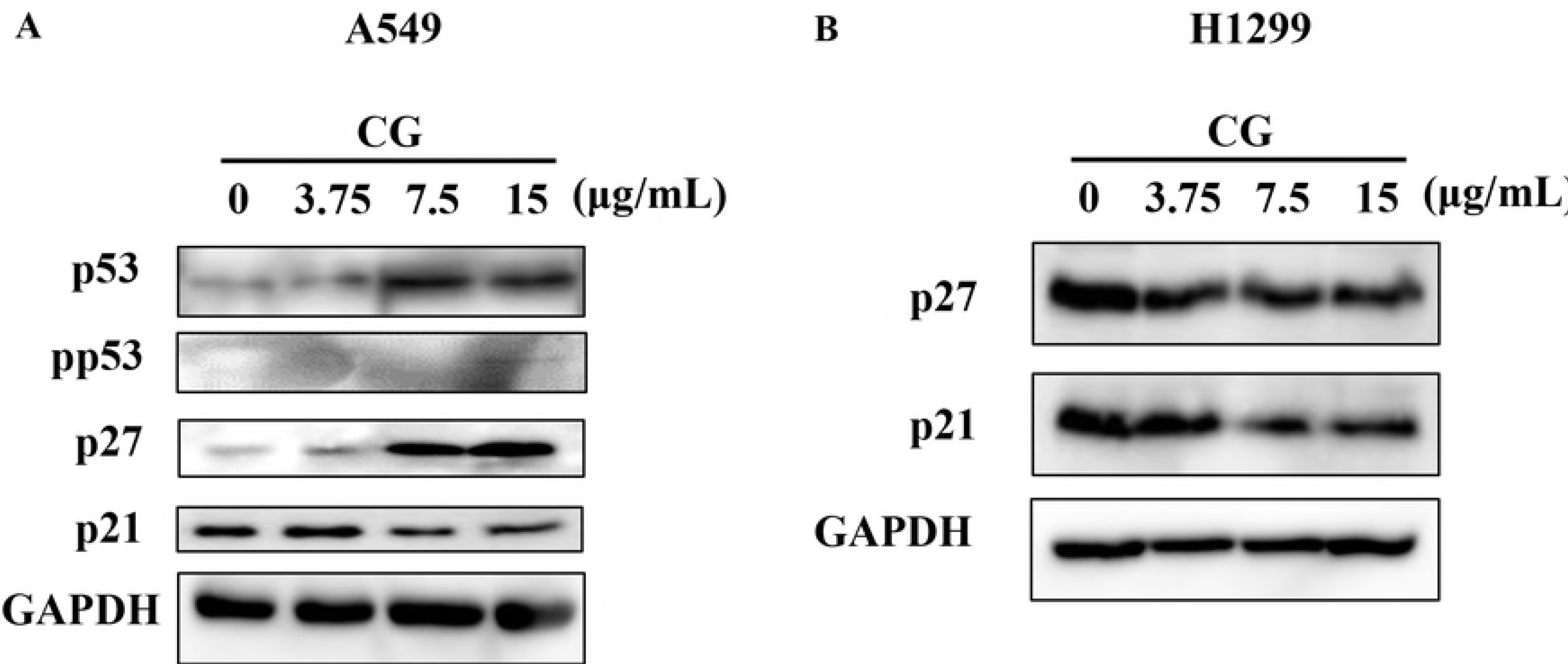
Effects of CG on p53-dependent apoptotic signaling in A549 cells and on the cell cycle phases in A549 and NCI-H1299 lung cancer cells. (A) Protein expression levels of p53, p27, p21, and GAPDH in A549 cells and (B) protein expression levels of p27, p21, and GAPDH in NCI-H1299 cells as determined by western blot analyses. A549 and NCI-H1299 cells were treated with various concentrations of CG for 48 h and compared with untreated cells. Cell cycle profiles of CG-treated A549 (C) and NCI-H1299 cells (D). The cells were treated with CG for 48 h, fixed, and stained with PI. A549 (E) and NCI-H1299 cells (F) in the sub-G1 phase. Results with **p < 0.005 were considered statistically significant. Protein expression levels of cyclin D1, cyclin E, cyclin A, and GAPDH in A549 (G) and NCI-H1299 cells (H) as determined by western blot analyses.

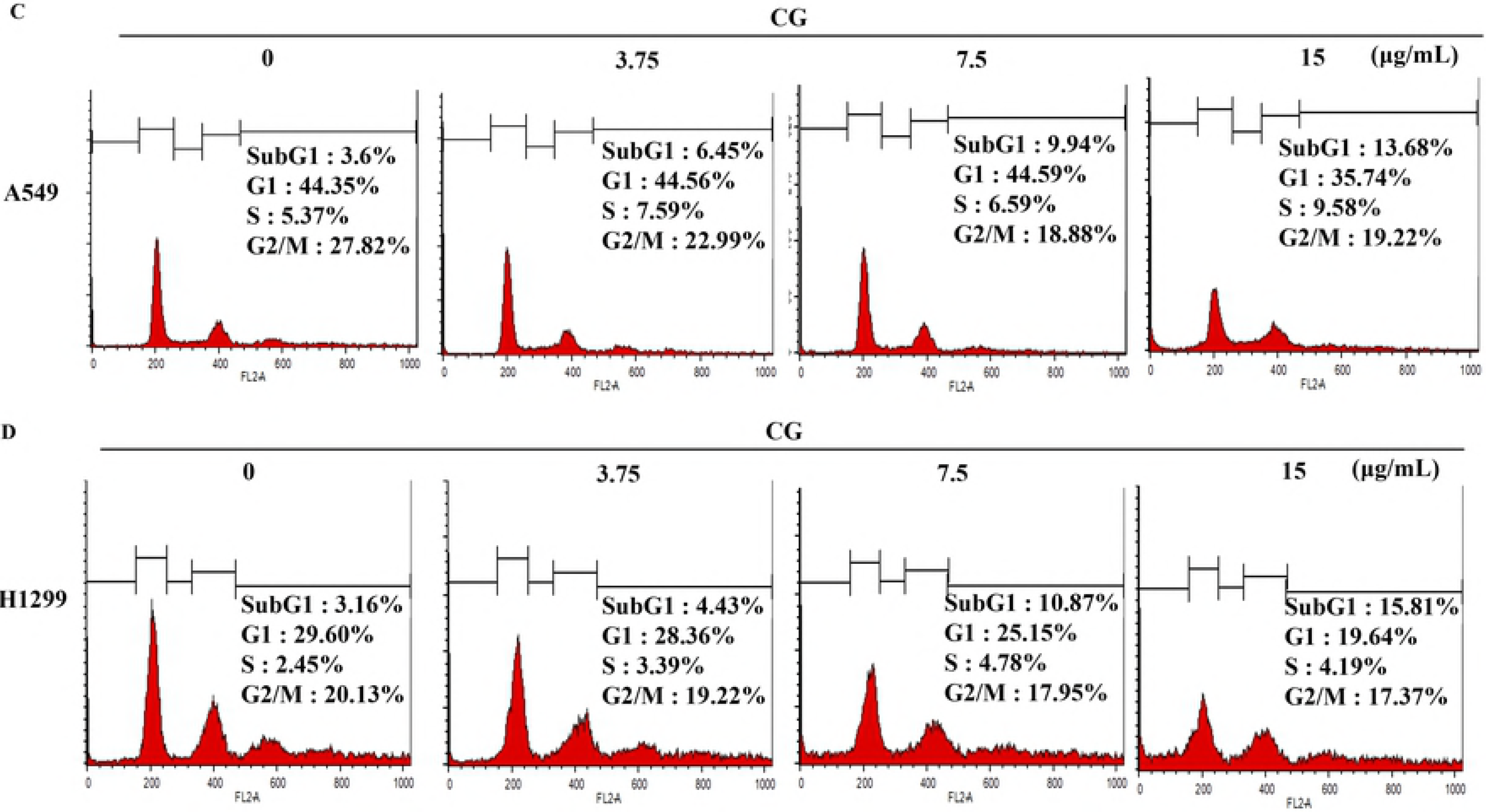

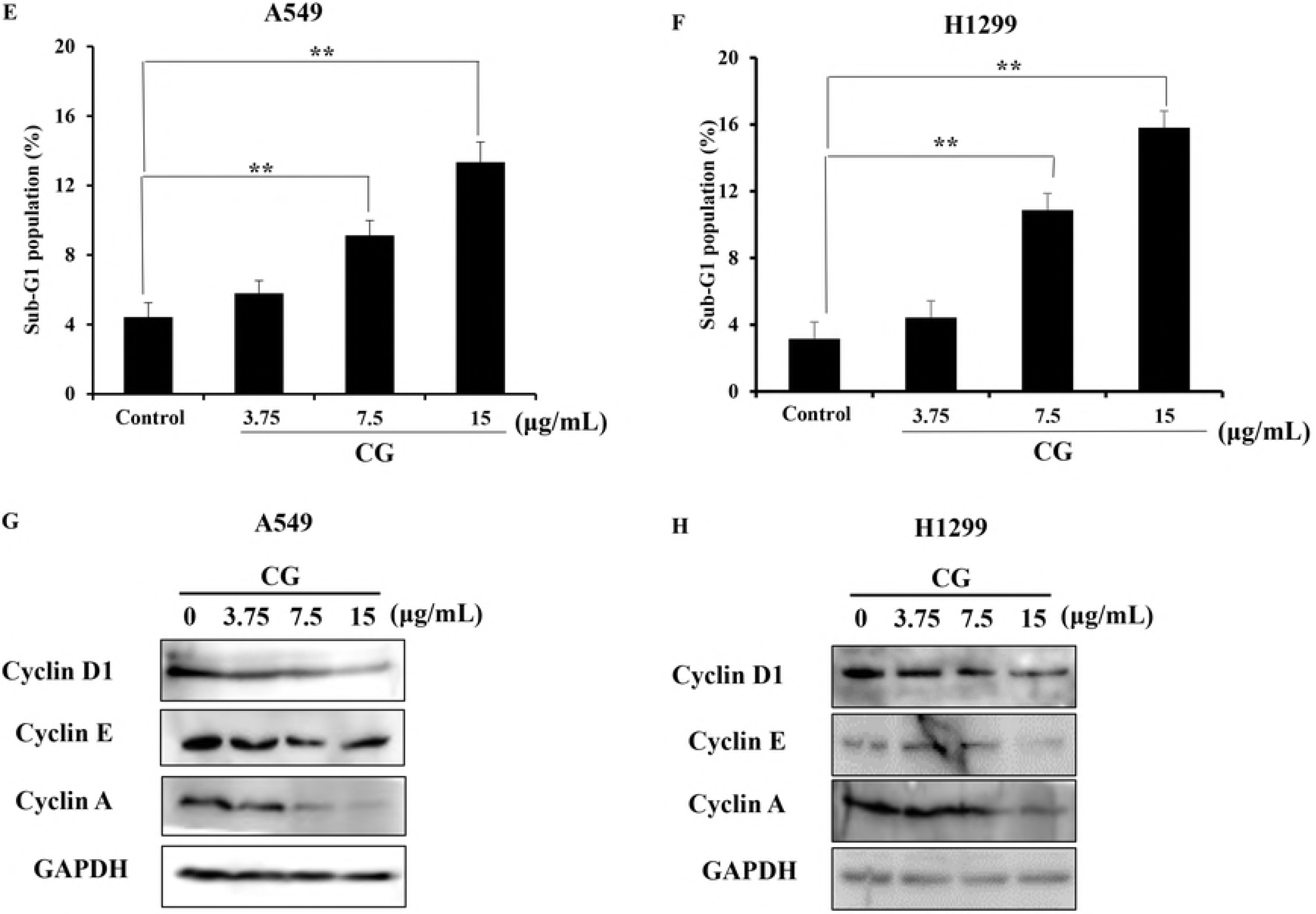

### CG induces the extrinsic pathway of apoptosis by activating death receptors and downregulating signaling

The extrinsic pathway of apoptosis is the other important pathway leading to apoptosis [25]. This pathway is distinct from the intrinsic pathway. The interactions between ligands and death receptors promote the formation of a death-inducing signaling complex (DISC), and caspase-8 is activated [26]. To confirm the mRNA levels of extrinsic pathway factors, real-time qPCR was performed. The results showed that factors in the extrinsic pathway tended to increase. The mRNA expression levels of death receptor 5 (DR5), FADD, FAS, and FASL were increased in CG-treated A549 (Fig. 5A) and NCI-H1299 cells (Fig. 5C). Furthermore, the pro-forms of caspase-8 expression levels decreased in a CG dose-dependent manner, and cleaved forms appeared following treatment with high concentrations of CG in A549 cells (Fig. 5B) and NCI-H1299 cells (Fig. 5D). These results demonstrated that CG was effective in inducing cell death through the process of apoptosis in A549 and NCI-H1299 lung cancer cells.

**Figure 5.**
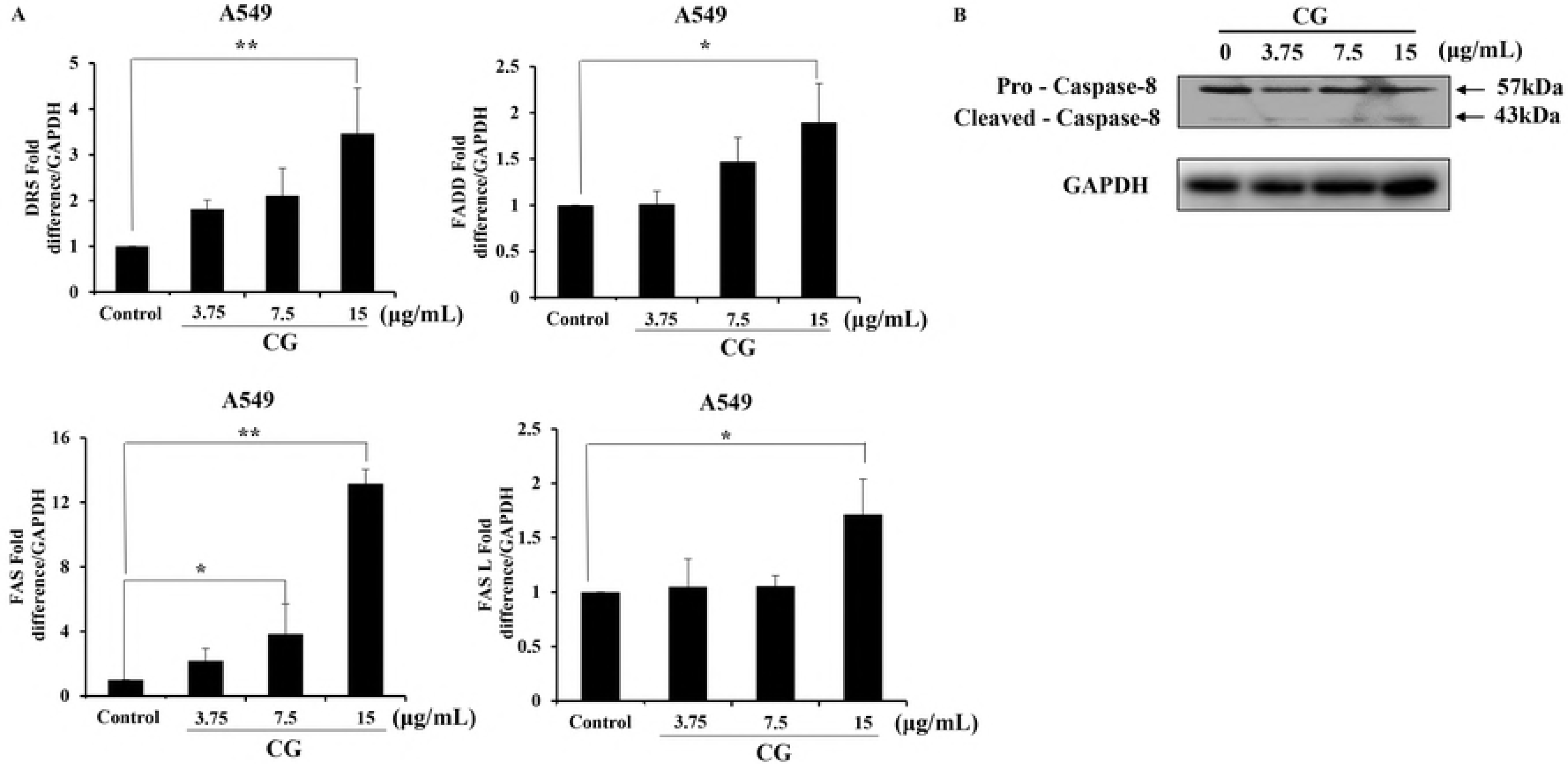
Effects of CG on extrinsic pathway-related factors in A549 and NCI-H1299 cells. mRNA levels of DR5, FADD, FAS, and FASL in A549 (A) and NCI-H1299 cells (C) as determined by qPCR. The graph was compiled from at least three replicates. Protein expression levels of the extrinsic pathway factors, pro-caspase-8 and its cleaved form, in A549 (B) and NCI-H1299 cells (D) as determined by western blot analyses. Cells were treated with various concentrations of CG for 48 h and compared with untreated cells. Results with *p < 0.05 and **p < 0.005 were considered statistically significant.

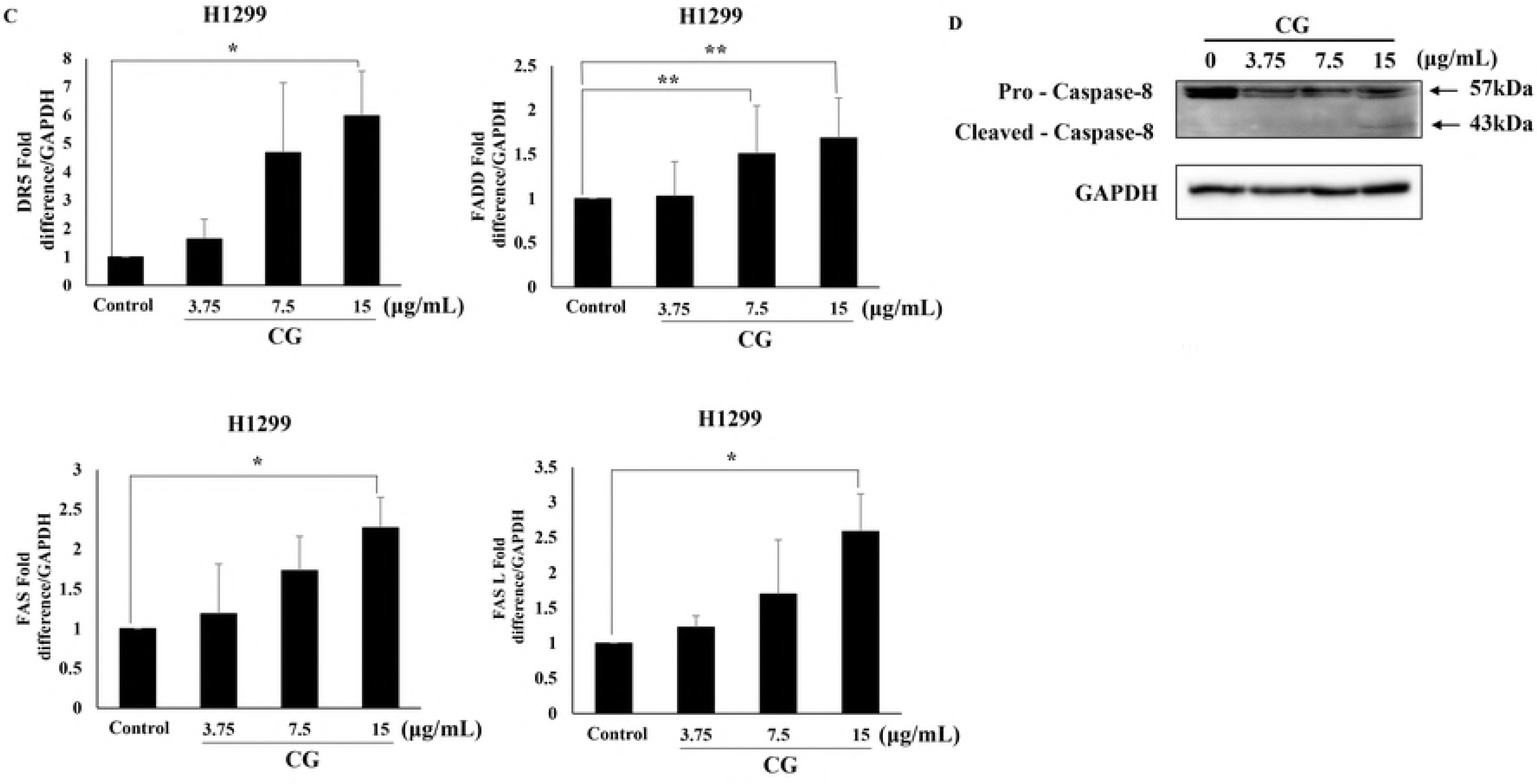

### CG has an apoptotic effect on mitochondrial and intracellular downregulation signals

The extrinsic and intrinsic pathways of apoptosis cross at the level of the mitochondria [27]. Activated caspase-8 cleaves the protein Bid into tBid, which is an activated form. This is mediated by members of the Bcl-2 family. tBid induces Bax-dependent outer mitochondrial membrane permeabilization and the release of cytochrome c [28]. Bid expression level was decreased, while Bax was enhanced in A549 cells following treatment with CG (Fig. 6A). Bcl-2, which is an inhibitory factor in the intrinsic pathway of apoptosis, also decreased, but the levels of Bcl-xL were not altered. These levels were similar in NCI-H1299 and A549 cells (Fig. 6E). Thus, Bax stimulated by Bid and Bcl-2 controlled MMP. The fluorescence of cells stained with JC-1 change from orange to green when apoptosis is in progress and MMP is decreasing. The orange fluorescence of A549 cells (Fig. 6B) and NCI-H1299 cells (Fig. 6F) exhibited a dose-dependent leftward shift following treatment with CG. Moreover, cytochrome c from mitochondrial membranes appeared in the cytosol in CG-treated A549 cells (Fig. 6C) and NCI-H1299 cells (Fig. 6G) as determined by western blot. Mitochondrion dysfunction is a very important signal in the intrinsic pathway of apoptosis [27], and mitochondria membrane collapse caused caspase-9 to be released. This study confirmed that factors such as caspase-9 and caspase-3, which are controlled by members of the Bcl-2 family (e.g., Bcl-2 and Bcl-xL) [29] decreased and were cleaved to induce apoptosis in a dose-dependent manner following CG treatment of A549 cells (Fig. 6D) and NCI-H1299 cells (Fig. 6H) as determined by western blot assay. The cleaved forms of these proteins were found following treatment at the highest concentration of CG in both cell lines. This suggests that these cleaved forms activated apoptosis. Finally, the key factor in programmed cell death and DNA repair, PARP, was cleaved by CG in the two types of cells (Fig. 6D and H). These results indicated CG-induced apoptosis following signaling in the intrinsic pathway.

**Figure 6.**
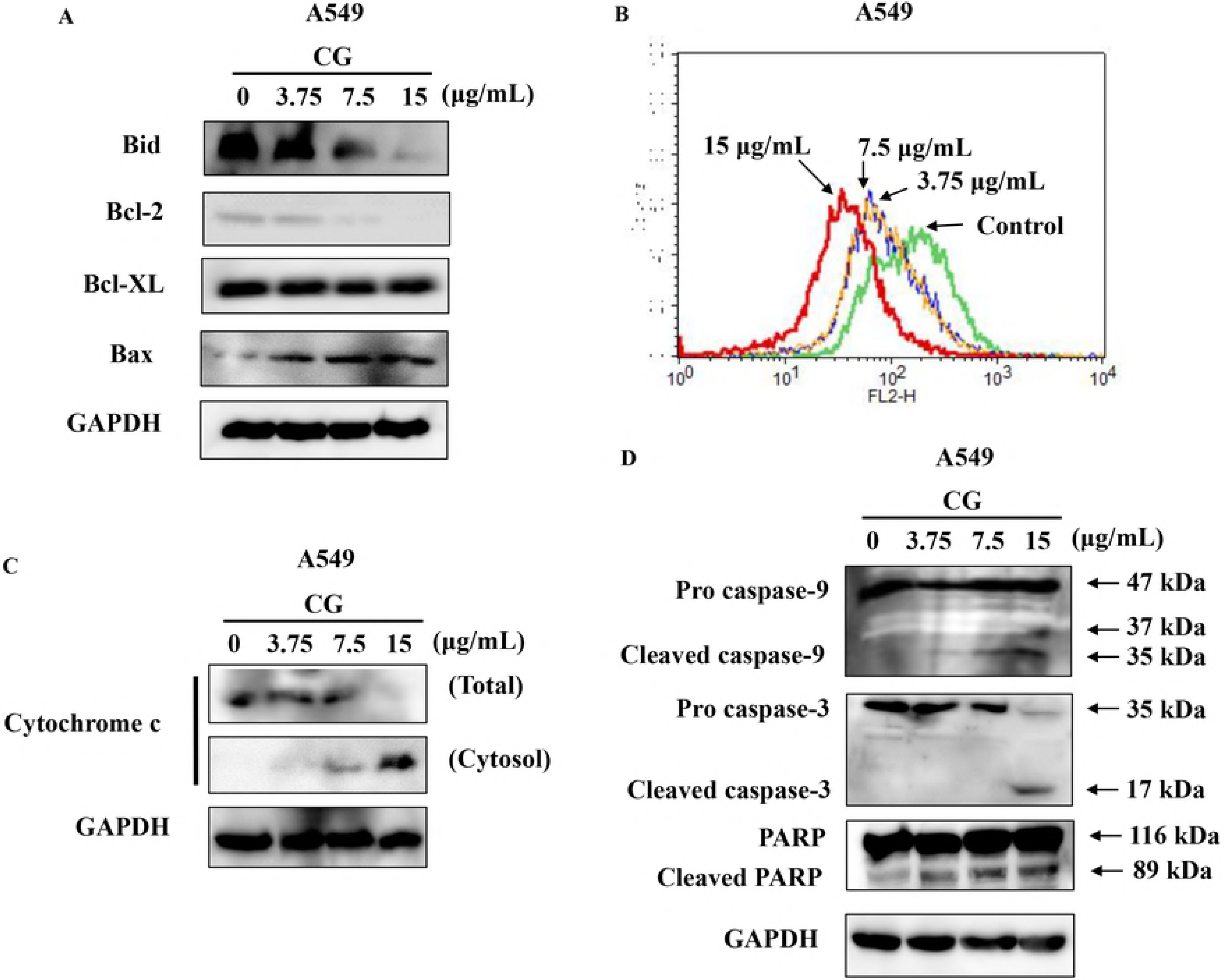
Effects of CG on MMP and intrinsic pathway-related factors in A549 and NCI-H1299 cells. Protein expression levels of BID, Bcl-2, Bcl-xL, BAX, and GAPDH in A549 (A) and NCI-H1299 cells (E) as determined by western blots. Cells were treated with various doses of CG extract for 48 h and compared with untreated cells. Histogram profiles of JC-1 aggregates (FL-2, orange) detected by flow cytometry of A549 (B) and NCI-H1299 cells (F). Western blots of cytochrome c protein levels in membranes and cytosol fractions and GAPDH in A549 (C) and NCI-H1299 cells (G). Cells were fractionated using an NE-PER nuclear cytoplasmic extraction reagent kit (Thermo, USA). Protein levels of the intrinsic pathway factors, caspase-9, caspase-3, PARP, and GAPDH in A549 (D) and NCI-H1299 cells (H) as determined by western blot analyses.

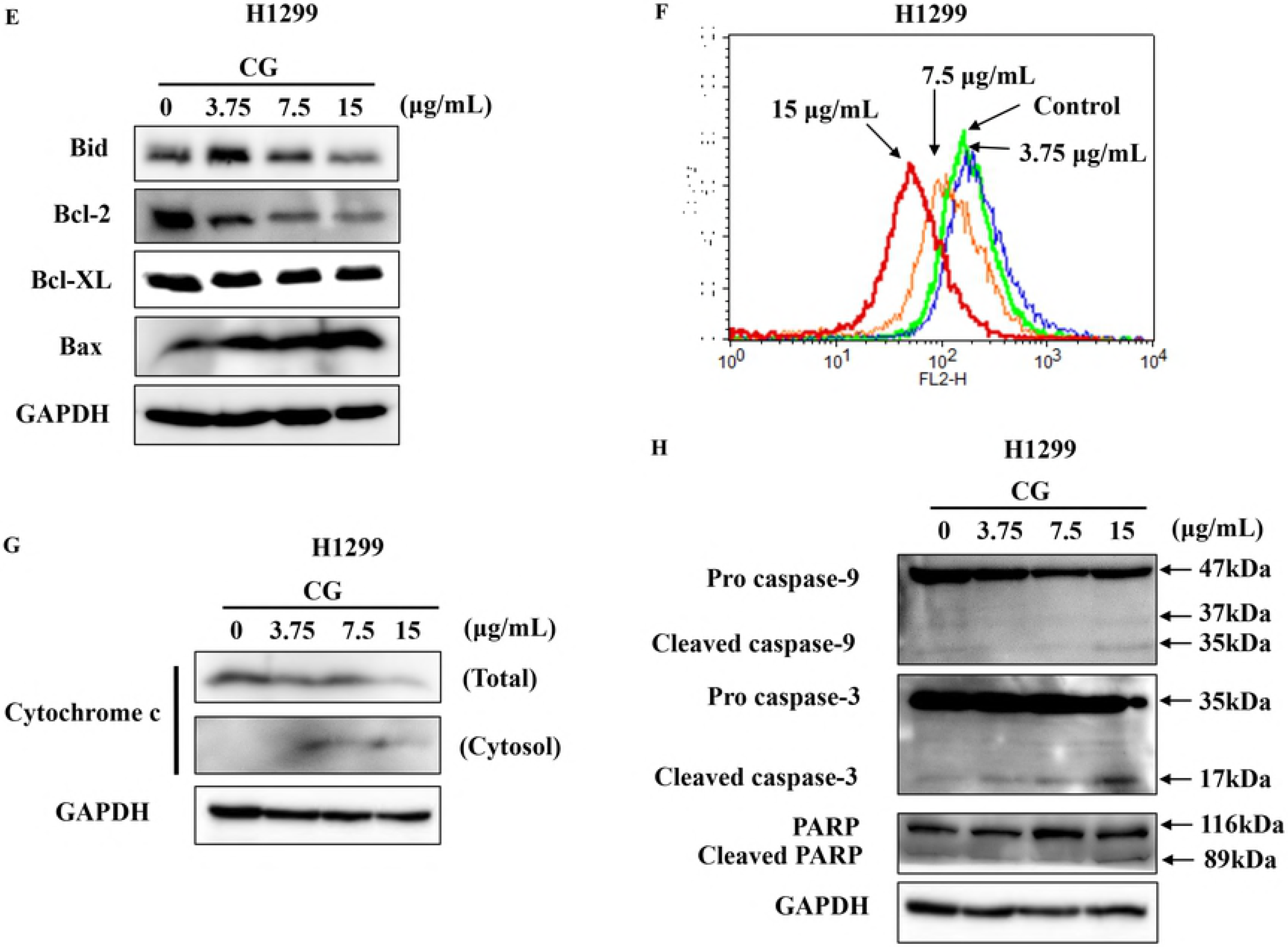

### CG generates ROS products

To determine how apoptosis occurs in p53-independent death of NCI-H1299 cells, we examined the generation of ROS, which is a main cause of apoptosis [23, 30]. There are many studies on ROS related to apoptosis. ROS levels can increase dramatically with environmental stress and result in significant damage called oxidative stress [5]. Therefore, we investigated the effect of CG on ROS production in A549 and NCI-H1299 cells. First, we examined ROS levels in A549 and NCI-H1299 cells treated with CG. After CG treatment for 48 h, these cells were stained with 2′,7′-dichlorofluorescin diacetate (DCF-DA), and the amount of ROS produced was determined by fluorescence microscopy. ROS accumulated dose dependently in A549 cells (Fig. 7A) and NCI-H1299 cells (Fig. 7C) following treatment with CG. Furthermore, ROS scavengers that had an anti-apoptotic role, superoxide dismutase 2 (SOD2) and catalase, decreased in A549 cells (Fig. 7B) and NCI-H1299 cells (Fig. 7D) in response to CG, but levels of the other ROS scavenger, thioredoxin (TXN), were unaltered (Fig. 7B and D). This study suggests that generated ROS might trigger or at least be implicated in CG-induced apoptosis.

**Figure 7.**
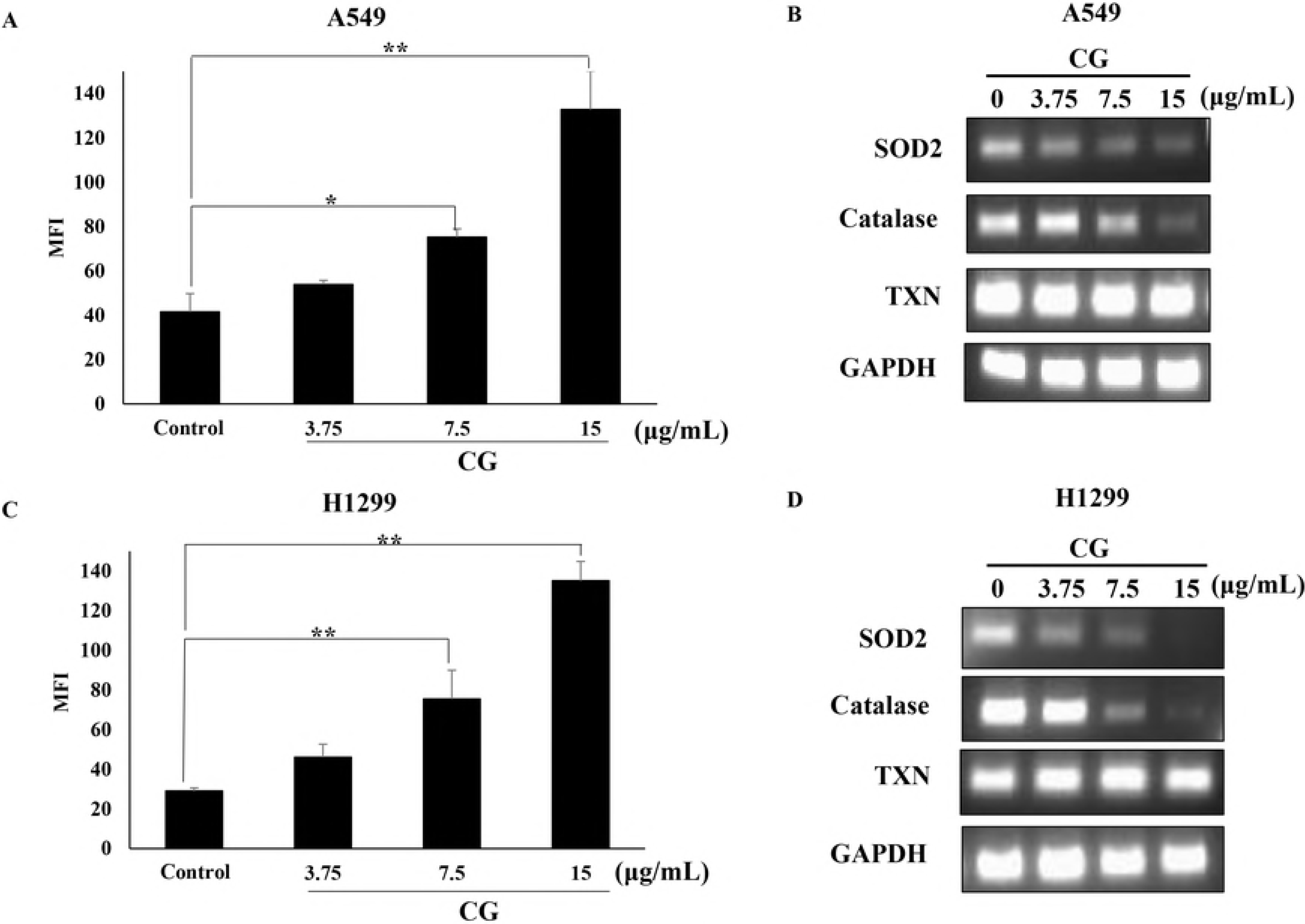
CG induced ROS generation in A549 and NCI-H1299 cells. CG treatment resulted in ROS generation in A549 (A) and NCI-H1299 cells (C). Cells were treated with CG for 48 h and examined using DCFDA fluorescence staining and a fluorescence microplate reader. The mRNA levels of superoxide dismutase (SOD2), catalase, thioredoxin (TXN), and GAPDH were determined by qPCR in A549 (B) and NCI-H1299 cells (D) treated with CG extract for 48 h. Results with *p < 0.05 and **p < 0.005 were considered statistically significant.

### ROS scavenger N-acetylcysteine (NAC) recovers cell viability

To confirm that CG extract induced apoptosis mediated by ROS generation, we used the ROS scavenger N-acetylcysteine (NAC) [23, 31] and examined cell viability and ROS generation. A549 and NCI-H1299 cells were pre-treated with NAC, and cell viability was confirmed. Cell viabilities dramatically recovered to over 100% in CG and NAC treatment groups, compared with the viabilities in the CG-treated groups of A549 cells (Fig. 8A) and NCI-H1299 cells (Fig. 8B). ROS generation levels also decreased with CG and NAC treatment of A549 (Fig. 8C) and NCI-H1299 cells (Fig. 8D), compared to the levels in CG-treated cells. Collectively, these results indicated that CG exerted anti-lung cancer effects through ROS-mediated apoptosis and the inhibition of ROS generation, which sufficiently blocked CG-induced apoptosis.

**Figure 8.**
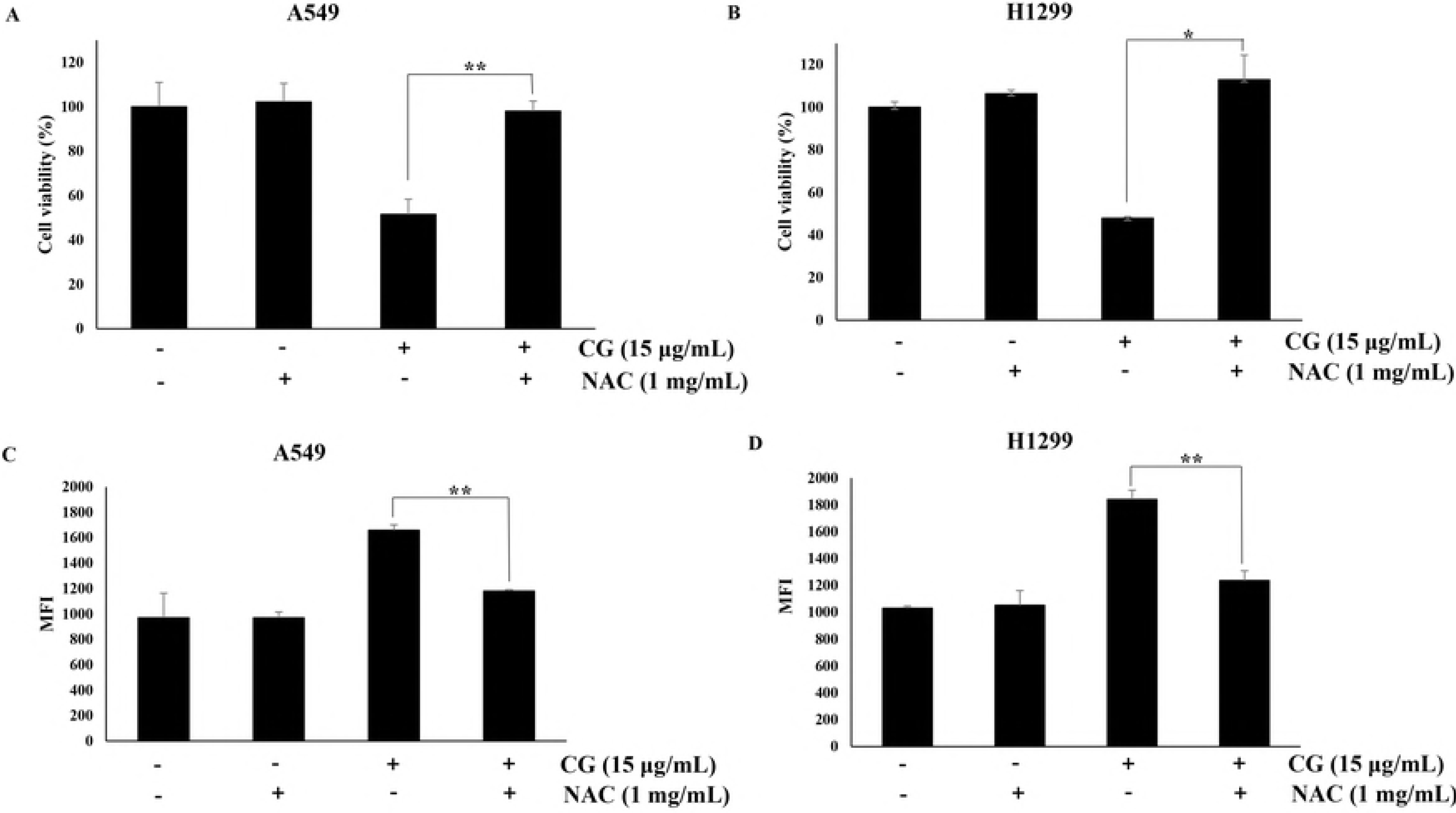
ROS scavenger N-acetylcysteine (NAC) recovered the cell viability and attenuated ROS production in CG-treated A549 and NCI-H1299 cells. Pre-treated NAC recovered the cell viability of A549 (D) and NCI-H1299 cells (B). A549 (C) and NCI-H1299 cells (D) were pretreated with NAC and then treated CG for 48 h. Then, the cells were stained with DCF-DA and examined using a fluorescence microplate reader. Results with *p < 0.05 and **p < 0.005 were considered statistically significant.

## Discussion

Many plant extracts are known as significant potential about anticancer effects [23, 32] such as inhibition of tumor growth, angiogenesis and metastasis. CG is one of plant extracts and few reports studied about the antiproliferative effects of the plant.

First, we analyzed that the chemical compounds of CG extract and the diverse rutinosides, well-known effective anti-cancer chemical [33], were detected. Among them, isorhamnetin-3-O-rutinoside which included high amount in CG extract has been investigated the apoptosis of human myelogenous erythroleukaemia cells [34], but had no effect about cytotoxicity in NSCLC cells (Fig. S1 in Supporting Information). However, CG extract has cytotoxic effects in non-small-cell lung cancer cells, especially in A549 (p53+/+) cells and NCI-H1299 (p53−/−) cells [35]. Not only cells’ morphology of the A549 cells and NCI-H1299 cells changed to more round shape and less cell-cell interaction than untreated cells as previous reported [36], but also the nuclear of both cells were fragmented and shrunk with CG treatment. Also, the late apoptosis was increased in a dose dependent CG treated A549 cells and NCI-H1299 cells. It meant that the cytotoxicity of these cells was because of apoptotic effects of CG extract.

p53 is well known as a tumor suppressor protein [22] and it stimulates the downstream factor, p27 [37]. p27 has ability to control the cell cycle which regulates cyclin D, the growth stimulation protein of the cyclin family [38]. Cyclin family such as cyclin D1, E, and A is involved in each specific phases of the cell cycle. During the cell cycle, hypodiploid fragmented DNA [21], one of apoptotic marker, is existed in Sub-G1 phage. In this study, p53 null NCI-H1299 cells were also used. As shown in Fig. 2B, the cytotoxicity of CG in NCI-H1299 cells was presented and it meant that CG extract induced cell death independent of p53 in NCI-H1299 cells. However, we confirmed that the expression levels of p53 and phospho-p53 (pp53) were increased in A549 cells (wild type p53+/+), treated with CG. As shown in Fig. 3E and 3F, the increasing rate of late apoptosis in NCI-H1299 cells was less than in A549 cells. These results demonstrated that p53 was strongly related to apoptosis, so p53 stimulated by CG in A549 cells induced a crucial apoptosis in compared to NCI-H1299 cells. Hence, p53 was one of essential factors to cause cytotoxic effect of CG in A549 cells. Also, p27 level was increase and controlled the cell cycle in A549 cells. During the cell cycle of the A549 cells and NCI-H1299 cells treated with CG, sub-G1 phase and S phage were increased, and G0/G1 phase population was decreased [23, 39]. It meant that the number of hypodiploid fragmented DNA in subG1 phage was increased and the cell cycle was limited by CG. Cyclin D1, a major player to activate G1 phase of cell cycle, was marginally inhibited and in S phage, cyclin A related to DNA replication [40] was decreased by CG in A549 cells and NCI-H1299 cells. In NCI-H1299 cells, cyclin E related to DNA replication, was also decreased by CG. Thus, it showed that CG extract caused adverse effects in the cell cycle of A549 cells and NCI-H1299 cells to stop the cells’ growth and induced apoptosis.

Apoptosis is elimination of damaged cells through ordered cell death system [6, 23]. There are two main pathways of apoptosis; extrinsic pathway and intrinsic pathway. We investigated that CG treatment increased the ratio of the death receptors, death ligands, adaptors and DISC of extrinsic pathway in A549 cells and NCI-H1299 cells. Also, extrinsic pathway and intrinsic pathway were connected through increased Bcl-2 and Bax by CG. Continuously, intrinsic programmed cell death by mitochondria outer membrane permeabilization (MOMP) [28] was induced and cytochrome c was released from mitochondria into the cytoplasm. This stimulation occurs downstream signaling cascade that cleaves other intrinsic caspases and PARP resulting in apoptosis [10, 23] by CG in both cells. CG extract induced apoptosis by stimulating the receptors and ligands of extrinsic pathway in A549 cells and NCI-H1299 cells. Additionally, the significant factors inducing the apoptotic intrinsic pathway in CG-treated A549 cells and NCI-H1299 cells were detected. It meant that CG induced apoptosis was depend on both pathways of intrinsic and extrinsic signaling in NSCLC.

There are numbers of evidences emphasizing the principal role of reactive oxidase species (ROS) production which induce cell death in various cancer cell types [13, 23]. Recent studies show that anticancer agents mediate their apoptotic effects through ROS and the ROS production is prevented by ROS scavenger blocking cell death [31]. In agreement with these studies, we observed CG-enhanced ROS generation in A549 and NCI-H1299 lung cancer cells in a dose dependent manner. Furthermore, the cell viabilities and ROS generation levels were recovered with the ROS scavenger, N-acetylcysteine (NAC), in A549 cells and NCI-H1299 cells.

In conclusion, CG has a cytotoxic effect against non-small lung cancer cells and inhibits via not only p53 dependent pathway in A549 cells but also p53 independent pathway in NCI-H1299 cells. This signaling is related with cell cycle through p27 in A549 cells. During cell cycle, the sub-G1 populations were increased with downregulation of cyclin D1 and cyclin A is decreased in A549 cells and NCI-H1299 cells. Also, cyclin E expression was controlled by CG in NCI-H1299 cells. Notably, CG induces the extrinsic and intrinsic apoptotic pathway mediated by death receptors, cytochrome c and caspases and inactivated by PARP through controlling Bcl-2 and Bax in A549 cells and H199 cells. CG also enhances ROS level and a ROS scavenger, NAC, recovers reduced cell viability by CG in A549 cells and NCI-H1299 cells. To summarize, CG plant extract has a profound anti-cancer effect and these experiments strongly support the apoptotic effects (Fig. 9). Through *in vivo* experiments in future study, the plants extract, CG, can be developed and used in lung cancer therapy.

**Figure 9.**
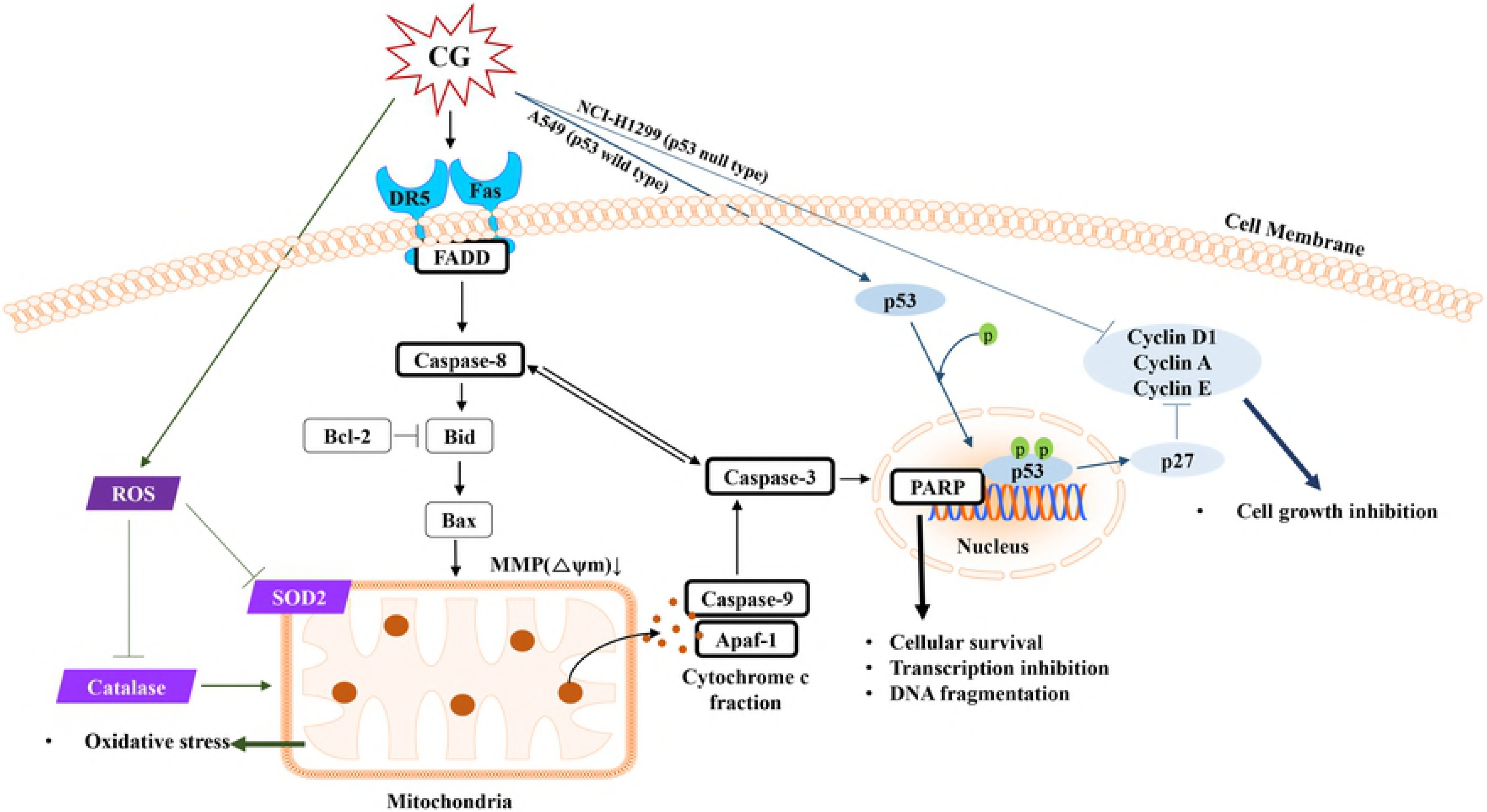
Schematic diagram illustrating CG-induced apoptotic effects in human NSCLC cell lines. CG triggers death receptor (DR5 and FAS)-and adaptor (FADD)-mediated apoptotic signaling pathways, as well as caspase-8 processing, resulting in cytochrome c release that is regulated by Bcl-2, Bid, and Bax. Subsequently, activated caspase-9 followed by caspase-3 cleaves PARP, resulting in triggering apoptosis. Moreover, CG induces apoptosis through ROS generation by controlling the ROS scavengers such as catalase and SOD2 in mitochondria. CG stimulates tumor suppressor p53 and cyclin-dependent kinase inhibitor p27. Cell cycle is also suppressed by a reduction in cyclin factors.

## Acknowledgements

This research was supported by Konkuk University in 2018.

## Supporting Information

**S1 Figure. Cell viability treated with Isohamnetin-3-O-rutinoside in the A549 and NCI-H1299 cells**. The viabilities of A549 and NCI-H1299 cells. A549 and NCI-H1299 cells were treated for 48 h with Isohamnetin-3-O-rutinoside. Viability was analyzed by MTS assay. The experiments were performed at least three times..

## Abbreviations used

CG: *Calotropis gigantea*
NSCLC: non-small cell lung cancer
FasL: Fas Ligand
FADD: Fas-associated protein with death domain
DR5: death receptor 5
PARP: poly (ADP-ribose) polymerase
ROS: reactive oxygen species

